# ipADMIXTURE: R package for inferring sub-population clusters based on genetic admixture

**DOI:** 10.1101/2020.03.21.001206

**Authors:** Chainarong Amornbunchornvej, Pongsakorn Wangkumhang, Sissades Tongsima

## Abstract

ipADMIXTURE is an R package to infer clusters and their phylogeny based on Q matrices of genetic admixture analysis. It is the first software of its kind to infer not just only clusters, but also the hierarchy of sub-populations w.r.t. the minimum number of ancestors that split any pair of clusters apart. Since inputs of the package, Q matrices, can be obtained from well-known software (ADMIXTURE, STRUCTURE, etc.) and the Q matrices are mandatory information that are used in genetic population structure study, our package has a potential to help scientists and researchers to find deeper explanation of admixture analysis in their studies. Our package comes with a user-friendly interface to make the software accessible for everyone.

## Required Metadata

### Current code version

**Table 1:** Code metadata (mandatory)

| Nr. | Code metadata description | Please fill in this column |
| --- | --- | --- |
| C1 | Current code version | v 0.1.0 |
| C2 | Permanent link to code/repository used for this code version | https://github.com/DarkEyes/ipADMIXTURE |
| C3 | Code Ocean compute capsule | — |
| C4 | Legal Code License | GPL-3 |
| C5 | Code versioning system used | git |
| C6 | Software code languages, tools, and services used | R, GitHub, TravisCI |
| C7 | Compilation requirements, operating environments & dependencies | 64-bit operating system, R (version $\geq 3.5.0$), R packages: stats, treemap, and ape |
| C8 | If available Link to developer documentation/manual | https://github.com/DarkEyes/ipADMIXTURE |
| C9 | Support email for questions |  |

## 1. Motivation and significance

Admixture analysis is one of fundamental methods in population biology for studying relativeness among populations as well as generic history [1]. It is the technique that tells us about histories of human migration [2], health study [3], rice population study [4] etc. Recently, high-throughput sequencing [5] and many technologies enable scientists to access big data of genetic sequences. With the overwhelming massive datasets, scientists face new challenges of knowledge discovery from massive data.

There are various methods that can perform data clustering on genetic data and report phylogeny of clusters, such as ipPCA [6], iNJclust [7], IP-CAPS [8], etc. However, no methods use admixture analysis that contains information of common ancestry to perform clustering. In population study, ADMIXTURE [9] and STRUCTURE [10] are widely used programs that report admixture analysis. Q matrices that contain admixture ratios from admixture are typically used to perform data clustering on some algorithm (e.g. k-means). However, no method uses admixture ratios to infer a hierarchy of sub-populations w.r.t. a minimum number of ancestors that split any pair clusters apart. Moreover, most data clustering algorithm provide clusters that are no biological interpretation in the aspect of admixture of ancestry.

With these reasons, we develop our R package, *ipADMIXTURE*. The goal of *ipADMIXTURE* is to make data clustering in population structure study more explainable. The reasons we use admixture analysis as a main core of our framework are as follows. First, individuals with similar clusters imply that they share the same mixture of ancestors. Second, using a phylogeny of sub-populations from admixture, we can tell a number of generations that two populations started splitting into different populations.

Our package is capable of:

- **Inferring individual assignments:** assigning individuals to be in the same cluster if they share similar admixture of ancestors; and
- **Estimating phylogeny of clusters:** estimating relativeness among clusters in a form of hierarchical tree; clusters that share a similar ancestor profile have small distance in the tree.

Given a huge number of R users in biology and related communities, we choose to implement our package in R to serve our target users. Our main goal is to provide an easy-to-use tool with more explainable results for researchers and practitioners.

In Section 2, we demonstrate how to use our package to infer clustering and phylogeny. Then, in Section 4, we show an example of the package application using the human 27 population dataset [11].

## 2. Software description

We provide details of ipADMIXTURE system architecture in Section 2.1, then we describe software functionality in Section 2.2.

### 2.1. Software Architecture

Our framework, ipADMIXTURE, has two different main components. The first component is for inferring clustering assignment w.r.t. a given Q matrix. The second component is for inferring a phylogeny tree of sub-clusters inferred from a first part.

We start with a first component that implemented in the ‘ipADMIXTURE function’. Given Q matrix as an input that contains admixture information, ipADMIXTURE uses top-down clustering scheme as a core of the framework. Figure 1 provides an overview of ipADMIXTURE software architecture. The binary clustering function is ‘biclustFunc’ function that deploys hierarchical clustering to perform clustering.

**Figure 1:**
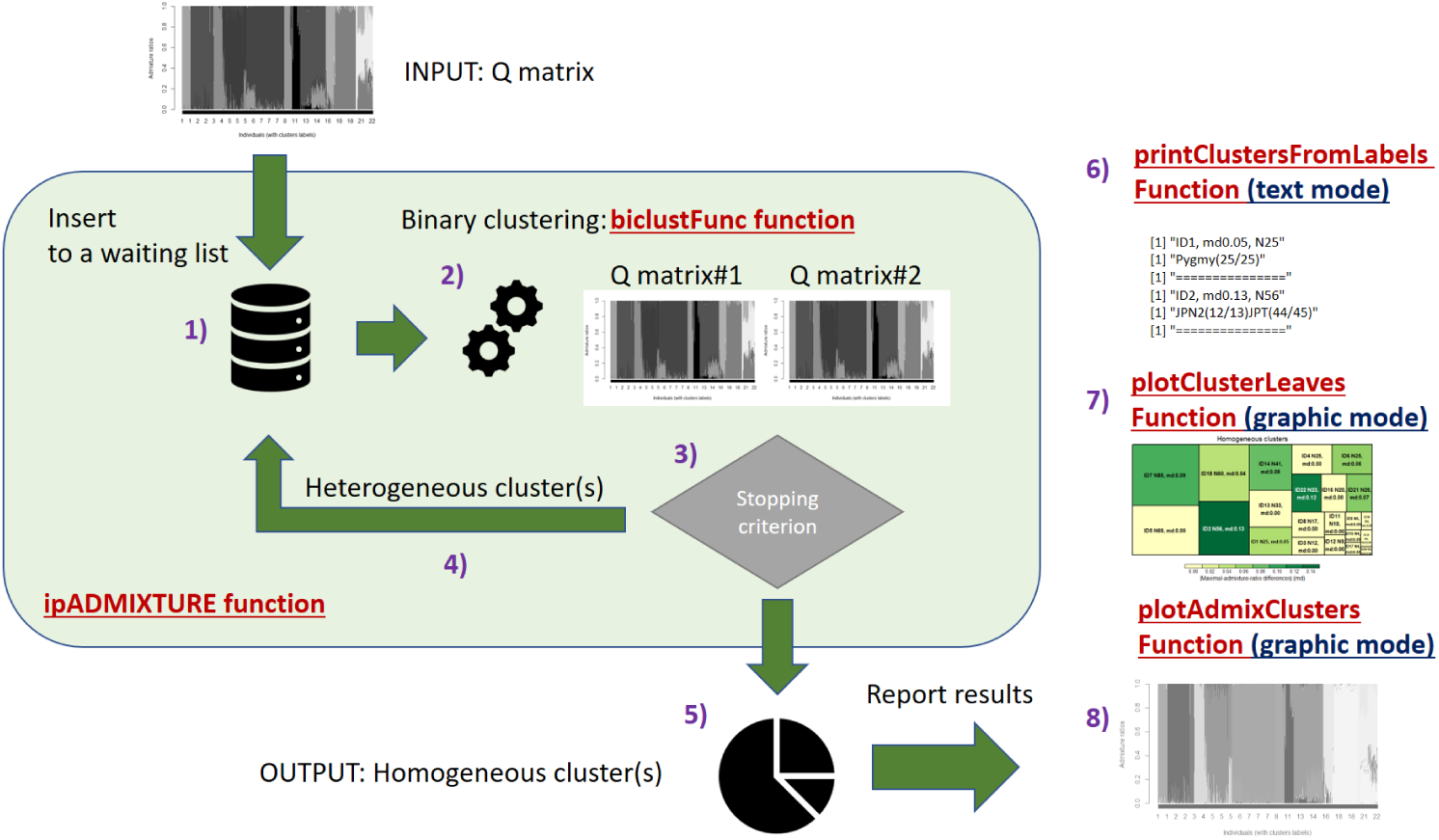
Software architecture of ipADMIXTURE in a part of data clustering. Given Q matrix as an input, ipADMIXTURE function provides clustering assignments and related visualization.

The top-down clustering scheme starts with a single cluster of all individuals, then 1) it inserts a cluster into a waiting list. Afterward, 2) the framework take some cluster *A* from a waiting list and divides a given cluster into two sub-clusters *A*1 and *A*2. Then, 3) the framework determines whether a given cluster *A* compares against its sub-clusters *A*1 and *A*2 is homogeneous w.r.t. a stopping criterion. If a given cluster *A* is not homogeneous, then 4) the frameworks puts sub-clusters *A*1 and *A*2 into a waiting-list. If a cluster *A* is homogeneous, then 5) it is assigned as an output cluster and nothing added into a waiting-list. In the next iteration, if anything is on the waiting list, the framework brings it to determine its homogeneity and the similar process continue until all individuals are assigned into homogeneous clusters. There are three ways of visualizing clustering assignment results in ipADMIXTURE, the details will be provided in Section 2.2. Shortly, the 6) ‘printClustersFromLabels’ function is used to print clustering results in text, 7) the ‘plotClusterLeaves’ function is used to plot degrees of homogeneity of clusters, and 8) the ‘plotAdmixClusters’ is used to plot admixture ratios with clusters.

For a phylogeny inference, given (1) a set of Q matrices with a number of ancestors varying from 2 to some constant *K*, and (2) a clustering assignment, the framework infer a phylogeny tree w.r.t. a minimum number of ancestors that split any pair clusters apart. Figure 2 provides details of software architecture in this part.

**Figure 2:**
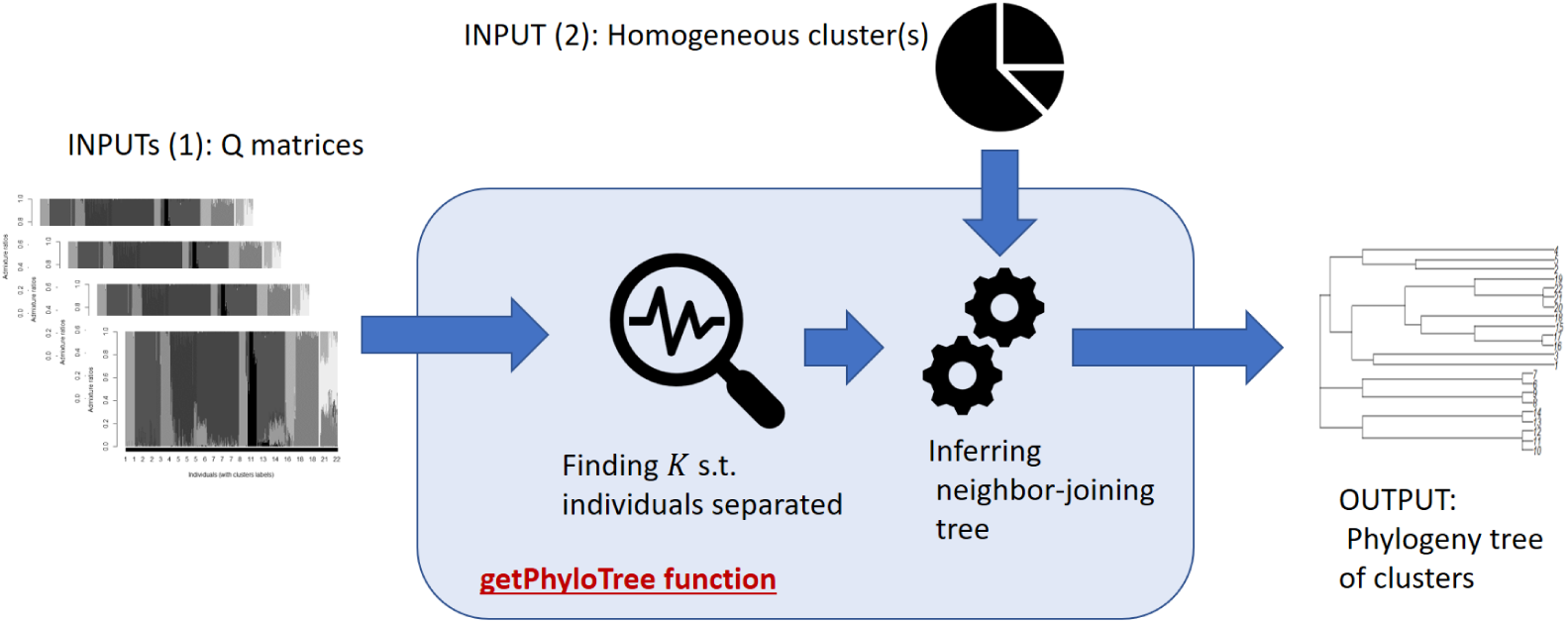
Software architecture of ipADMIXTURE in a part of phylogeny inference.

For a hierarchical clustering, we use an R *stats* package [12]. For phylogeny inference, we use a neighbor-joining method [13] in *ape* package [14]. We use *treemap* package [15] for visualizing homogeneous clusters and *graphics* package [12] for plotting admixture ratios w.r.t clusters.

### 2.2. Software Functionalities

The main tasks of ipADMIXTURE are 1) inferring clusters based on ancestry profiles; individuals who share the same ancestors belong to the same cluster, and 2) inferring a phylogeny tree of clusters; clusters that share similar ancestors stay closer in a tree.

#### 2.2.1. Clustering and homogeneity of clusters

We use hierarchical clustering as a main binary clustering engine. We use a “Maximum of magnitude-difference of admixture ratios”. Given *Q* is a Q matrix of cluster *C* where *Q*(*i, j*) represents a value of admixture ratio of *j*th ancestor for individual *i*, clusters *C*_1_ and *C*_2_ that are sub-clusters of *C* where *C* = *C*_1_ ∪ *C*_2_. The “Maximum of magnitude-difference of admixture ratios” is defined as follows.

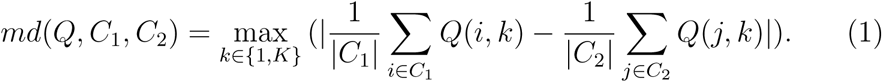

Where *md*(*Q, C*_1_, *C*_2_) ∈ [0, 1]. If *md*(*Q, C*_1_, *C*_2_) is high, then *C*_1_ and *C*_2_ has a high degree of different ancestry profile, which implies that *C*_1_ and *C*_2_ should be split into different clusters. In contrast, if *md*(*Q, C*_1_, *C*_2_) is low, then it implies that *C*_1_ and *C*_2_ have similar ancestry profile. Hence, *C*_1_ and *C*_2_ belong to the same cluster and *C* is a homogeneous cluster. In our framework, users can set a threshold *λ* ∈ [0, 1] to define a degree of clustering homogeneity. For any cluster, if its *md* < *λ*, then a cluster is a homogeneous cluster.

#### 2.2.2. Phylogeny inference

To infer a phylogeny tree of homogeneous clusters, we use a set of Q matrices with various numbers of ancestors and a set of homogeneous clusters as inputs. We report a phylogeny tree that represents relativeness. between clusters an an output.

In the first step, for any Q matrix of *k* ancestors, the framework assigns individuals to belong to *k* ancestry clusters w.r.t. a value of maximum admixture ratio. For any individual *i*, it belongs to *j*th ancestry cluster if *i* has a value of maximum admixture ratio that belongs to *j*th ancestor. Then, after we identify ancestry clusters for all Q matrices, we perform the next step.

For any pair of homogeneous clusters *A, B* and from all Q matrices, we find a minimum ancestor number *k*\* s.t. the majority of individuals between two clusters belong to different ancestry clusters. A similarity value between *A* and *B* is *k*\*. We create a distance matrix *D* between all pairs of homogeneous clusters based on *k*\*, then the framework constructs a phylogeny tree of these homogeneous clusters using Neighbor-Joining method (NJ) [13], which is one of the well-known approaches to infer phylogenetic trees.

NJ method constructs a phylogeny tree based on genetic distances. A length of edge in a tree created by NJ method represents a genetic distance between individuals. In this work, we use *D* as our genetic distance matrix to construct NJ phylogeny tree of clusters. Hence, a path length between clusters within a phylogeny tree represents a degree of difference of ancestry profiles between clusters.

## 3. Comparison to existing iterative-pruning clustering methods on genetic data

In this section, we compare the feature of ipADMIXTURE to other existing iterative pruning methods, including ipPCA, iNJclust and IPCAPS. The features are listed in Table 2, both ipADMIXTURE and IPCAPS are under GPL-3 license while ipPCA and iNJclust are unidentified. ipPCA was implemented using MATLAB while other tools were developed using non-comercial languages e.g. C++ for iNJclust and R for ipADMIXTURE and IPCAPS.

**Table 2:** Comparison of ipADMIXTURE to existing iterative pruning methods.

| Package | Source | Language | Available functions | License |
| --- | --- | --- | --- | --- |
| ipPCA | [6] | MATLAB | Clusters and their relativeness on PCA | - |
| iNJclust | [7] | C++ | Clusters and their relativeness on ASD | - |
| IPCAPS | [8] | R | Clusters and their relativeness on PCA | GPL-3 |
| ipADMIXTURE |  | R | Clusters and their relativeness on ancestry profiles | GPL-3 |

While all tools offer genetic clustering, ipADMIXTURE is the only one tool that clusters individuals based on their ancestry profile. ipPCA and IPCAPS clusters individuals based on the PCA-space distance and iNJclust produces Neighbor-Joining clusters using allele-sharing distance (ASD). Morever, ipADMIXTURE offers intuitive visualizations including Q matrices sorted by resulting clusters, phylogeny and homogeneity plot under the R ecosystem.

## 4. Case-study: inferring relativeness of human 27 populations

In this example, we have data set of human 27 population data publised by Xing, J., et al. [11]. The dataset consists of 544 individuals from 27 populations. The Q matrices from this data are provided in this package. The following steps are the simple way to use our package. First, we run the ipADMIXTURE using the Q matrix of huuman 27 population dataset (human27pop Qmat[[11]]) where the number of ancestors K =12 and the homogeneous threshold *λ* = 0.15.

~~~
**R**>**library** (ipADMIXTURE)
**R**>h27pop obj**<** −ipADMIXTURE(Qmat=human27pop_Qmat [[1 1 ]]
   , admixRatioThs =0.15)
~~~

The h27pop obj is an object of ipADMIXTURE class. It contains results of clustering. To show the result of clustering in admixture ratios, we can use the following command.

~~~
**R**>plotAdmixClusters (h27pop_obj)
~~~

The result of admixture plot is at Figure 3. Here, we have 22 clusters with 12 ancestors.

**Figure 3:**
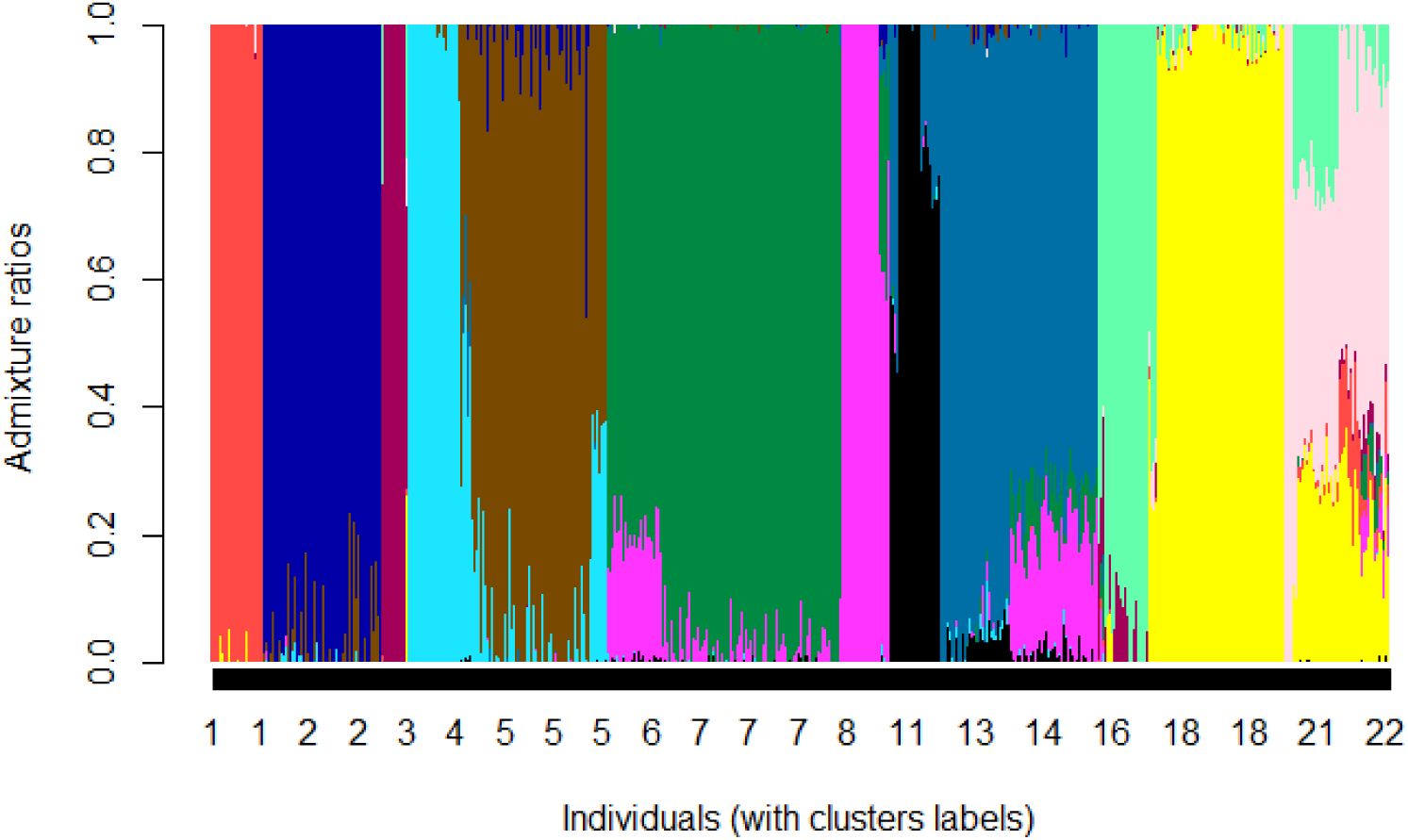
Admixture ratios with clusters. The horizontal axis represents each individual with its cluster ID. The vertical axis represents values of admixture ratios in colors. Each color represents each ancestor. Individuals who share the same ancestry profile have similar pattern of colors in this plot. Since we have 12 ancestors, there are 12 colors in the plot.

We also can show a result of clustering in the plot by focusing on levels of homogeneity and number of individuals within clusters using the following command.

~~~
**R**>plotClusterLeave (h27pop_obj)
~~~

The result of this plot is in Figure 4. Squares are clusters. A cluster with a larger number of members have a larger square. A color represents a degree of cluster homogeneity. Lighter color implies higher degree of homogeneity.

**Figure 4:**
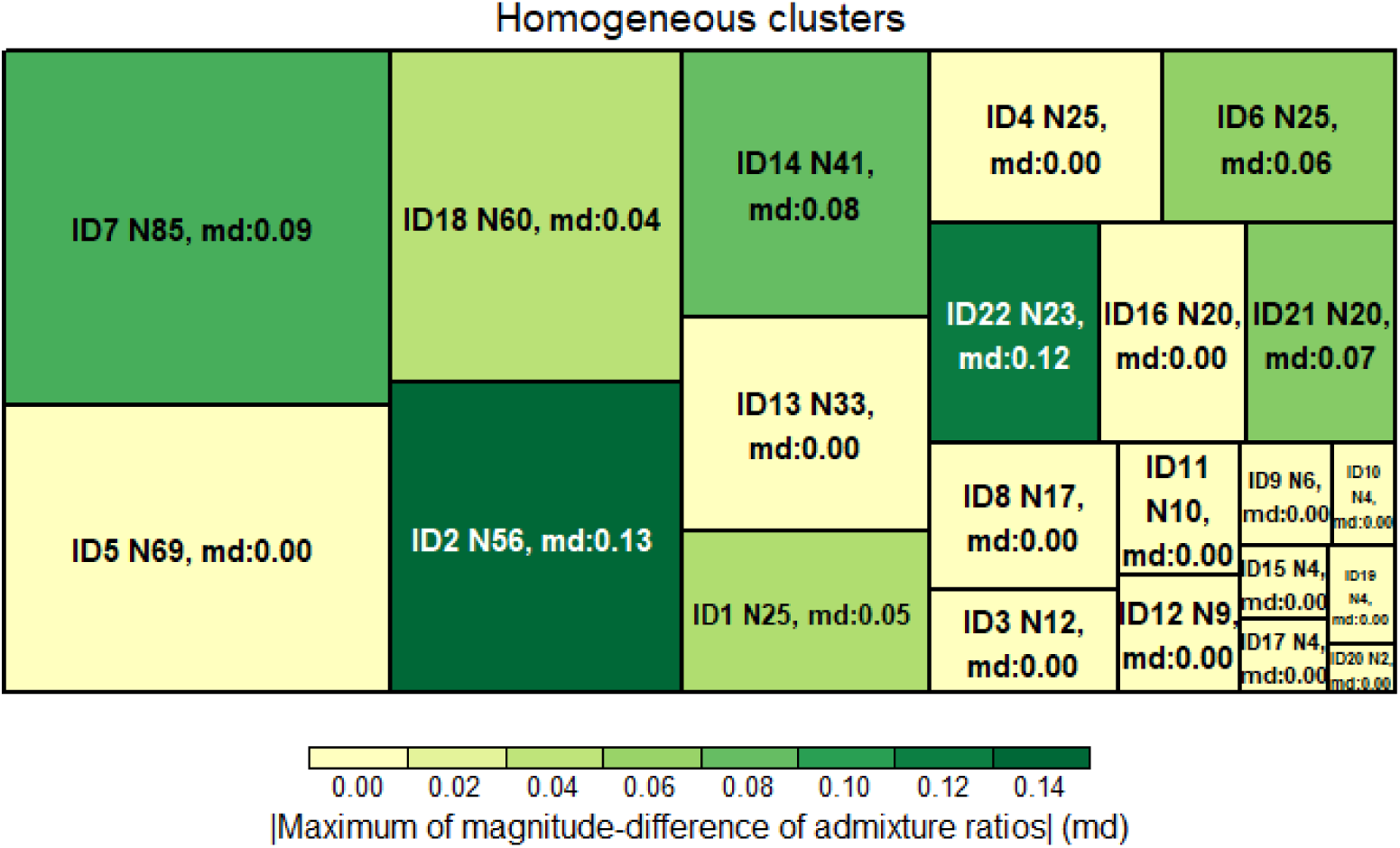
The clustering result with levels of homogeneity and number of individuals within clusters. Squares represent clusters. In each square, there are cluster ID, the number of members within a cluster (N) and the homogeneity degree in Equation 1 (md). A size of square represents a number of cluster members and the color represents a value of md. A cluste with higher degree of homogeneity has a lighter color.

Next, given a list of Q matrices from *K* = 2 to *K* = 12 (human27pop Qmat), and a clustering assignment vector (h27pop obj$indexClsVec), we infer a phylogeny tree of clusters using the following command.

~~~
**R**>out**<**−getPhyloTree(human27pop_Qmat, h27pop_obj**$**indexClsVec)
**R**>**plot** (out**$**tree)
~~~

The result of phylogeny inference is at Figure 5. In the figure, each leave node represents a cluster ID and a path length represents how far between two cluster are. According to the result, our framework correctly identified the structure human populations where people from the same Continent belong to the same sub-tree.

**Figure 5:**
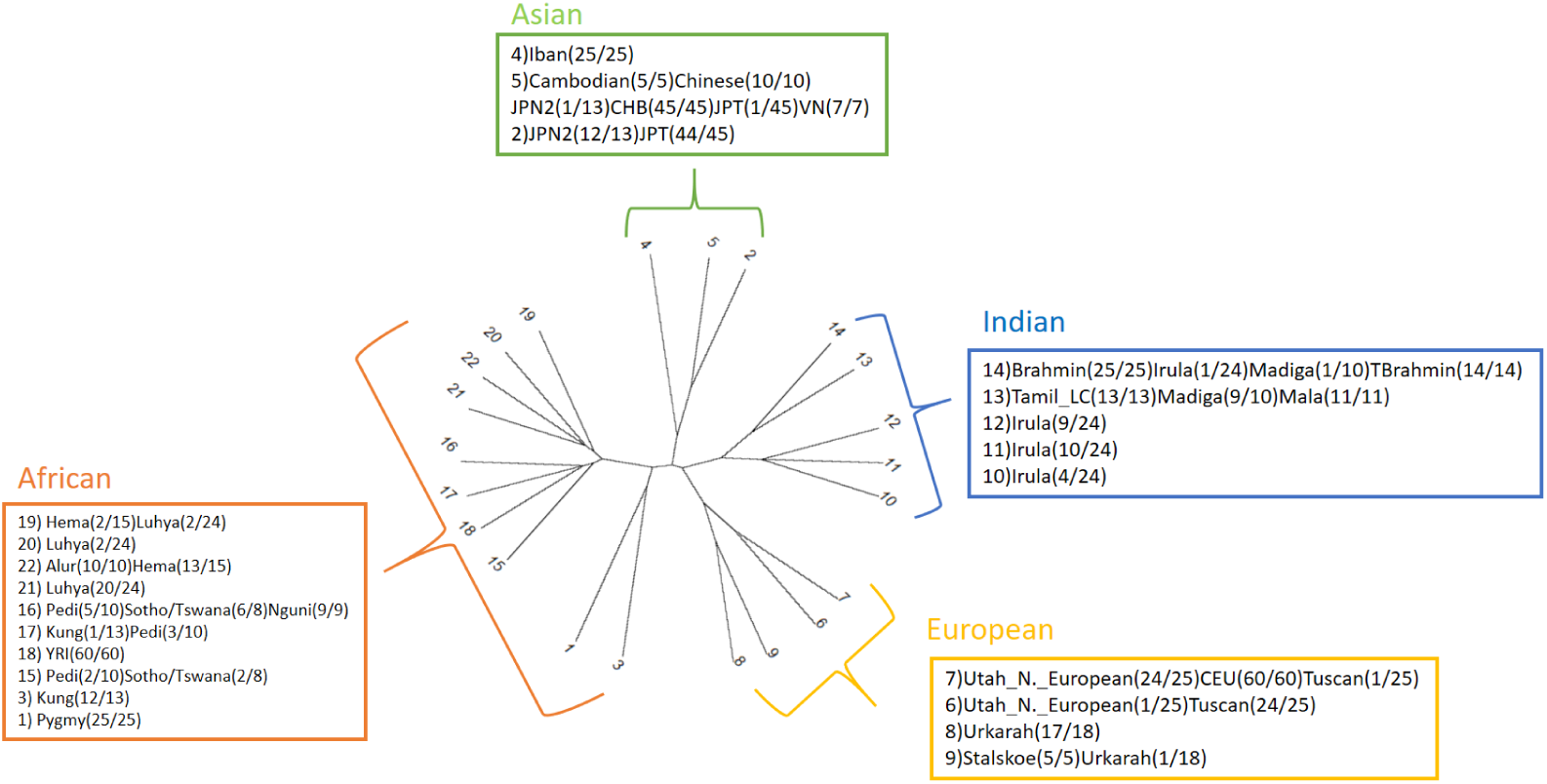
An inferred phylogeny tree from our framework using Q matrices from 27 populations dataset. Leave nodes represent clusters attached with cluster IDs. Square boxes contain clusters following their geography continents. Our framework correctly identified the structure human populations where people from the same Continent belong to the same sub-tree.

## 5. Conclusions

In this work, we presented a software package in R, ipADMIXTURE, for genotypic data clustering based on individual ancestry profile of admixture analysis. Clusters inferred by ipADMIXTURE have biological meaning; individuals share the same cluster have the same ancestry profile. Our framework is the first to use ancestry profiles of admixture analysis to perform iterative pruning clustering. Not just only reporting clusters, ipADMIXTURE also reports a phylogeny of clusters based on how far ancestry profiles between clusters. Taking inputs from any well-known software of admixture analysis (STRUCTURE or ADMIXTURE), our framework is reliable and has no limitation of numbers of SNPs and individuals it can perform analysis. The framework has a potential to support biologists, researchers, or practitioners to find their genotypic clusters that have biological interpretation w.r.t. ancestry profiles.

## 6. Conflict of Interest

We wish to confirm that there are no known conflicts of interest associated with this publication and there has been no significant financial support for this work that could have influenced its outcome.

## Current executable software version

**Table 3:** Software metadata (optional)

| Nr. | (Executable) software meta-data description | Please fill in this column |
| --- | --- | --- |
| S1 | Current software version | v 0.1.0 |
| S2 | Permanent link to executables of this version | https://github.com/DarkEyes/ipADMIXTURE |
| S3 | Legal Software License | GPL-3 |
| S4 | Computing platforms/Operating Systems | Linux, OS X, Microsoft Windows, Unix-like |
| S5 | Installation requirements & dependencies | 64-bit operating system, R (version $\geq 3.5.0$), R packages: stats, treemap, and ape |
| S6 | If available, link to user manual - if formally published include a reference to the publication in the reference list | https://github.com/DarkEyes/ipADMIXTURE |
| S7 | Support email for questions |  |

## Notes

https://github.com/DarkEyes/ipADMIXTURE

## References

[1] B. M. Peter, Admixture, population structure, and f-statistics, Genetics 202 (4) (2016) 1485–1501.

[2] G. Hellenthal, G. B. Busby, G. Band, J. F. Wilson, C. Capelli, D. Falush, S. Myers, A genetic atlas of human admixture history, Science 343 (6172) (2014) 747–751.

[3] A. P. Reiner, E. Ziv, D. L. Lind, C. M. Nievergelt, N. J. Schork, S. R. Cummings, A. Phong, E. G. Burchard, T. B. Harris, B. M. Psaty, et al., Population structure, admixture, and aging-related phenotypes in african american adults: the cardiovascular health study, The American Journal of Human Genetics 76 (3) (2005) 463–477.

[4] S. R. McCouch, M. H. Wright, C.-W. Tung, L. G. Maron, K. L. McNally, M. Fitzgerald, N. Singh, G. DeClerck, F. Agosto-Perez, P. Korniliev, et al., Open access resources for genome-wide association mapping in rice, Nature communications 7 (1) (2016) 1–14.

[5] D. M. Blei, P. Smyth, Science and data science, Proceedings of the National Academy of Sciences 114 (33) (2017) 8689–8692. arXiv:https://www.pnas.org/content/114/33/8689.full.pdf, doi: 10.1073/pnas.1702076114. URL https://www.pnas.org/content/114/33/8689

[6] A. Intarapanich, P. J. Shaw, A. Assawamakin, P. Wangkumhang, C. Ngamphiw, K. Chaichoompu, J. Piriyapongsa, S. Tongsima, Iterative pruning pca improves resolution of highly structured populations, BMC bioinformatics 10 (1) (2009) 382.

[7] T. Limpiti, C. Amornbunchornvej, A. Intarapanich, A. Assawamakin, S. Tongsima, injclust: iterative neighbor-joining tree clustering framework for inferring population structure, IEEE/ACM transactions on computational biology and bioinformatics 11 (5) (2014) 903–914.

[8] K. Chaichoompu, F. Abegaz, S. Tongsima, P. J. Shaw, A. Sakuntabhai, L. Pereira, K. Van Steen, Ipcaps: an r package for iterative pruning to capture population structure, Source code for biology and medicine 14 (1) (2019) 2.

[9] D. H. Alexander, J. Novembre, K. Lange, Fast model-based estimation of ancestry in unrelated individuals, Genome research 19 (9) (2009) 1655–1664.

[10] M. J. Hubisz, D. Falush, M. Stephens, J. K. Pritchard, Inferring weak population structure with the assistance of sample group information, Molecular ecology resources 9 (5) (2009) 1322–1332.

[11] J. Xing, W. S. Watkins, D. J. Witherspoon, Y. Zhang, S. L. Guthery, R. Thara, B. J. Mowry, K. Bulayeva, R. B. Weiss, L. B. Jorde, Fine-scaled human genetic structure revealed by snp microarrays, Genome research 19 (5) (2009) 815–825.

[12] R Core Team, R: A Language and Environment for Statistical Computing, R Foundation for Statistical Computing, Vienna, Austria (2019). URL https://www.R-project.org/

[13] N. Saitou, M. Nei, The neighbor-joining method: a new method for reconstructing phylogenetic trees., Molecular biology and evolution 4 (4) (1987) 406–425.

[14] E. Paradis, K. Schliep, ape 5.0: an environment for modern phylogenetics and evolutionary analyses in R, Bioinformatics 35 (2018) 526–528.

[15] M. Tennekes, treemap: Treemap Visualization, r package version 2.4-2 (2017). URL https://CRAN.R-project.org/package=treemap

